# Biomanufacturing of a functional microbial phytase in an insect host

**DOI:** 10.1101/2024.09.07.611833

**Authors:** C. Retief, S. Kumar, K. Tepper, M. Maselko

## Abstract

**Background:** Insects, such as Black Soldier Flies (*Hermetia illucens*), are increasingly used as sustainable animal feed ingredients that can be reared on plentiful organic substrates such as agricultural residues and pre-consumer food waste. Genetically engineering insects to heterologously express feed additive enzymes has the potential to generate more value from organic waste, while improving livestock health and productivity.

Phytases are widely used feed additive enzymes that hydrolyse the phosphate groups from the myo-inositol backbone of phytic acid, a phosphate rich antinutrient compound that monogastric animals cannot efficiently digest. Dietary phytase supplementation improves absorption of phosphorous, proteins, and cationic nutrients, while mitigating the negative environmental effects of phytic acid rich excreta.

**Results:** We evaluated the potential of using insects to biomanufacture microbial feed additive enzymes by engineering the model insect, *Drosophila melanogaster*, to express phytases. One histidine acid phytase, three beta propellor phytases, three purple acid phosphatases, and one PTP-like phytase were selected for screening in *D. melanogaster*. Transgenic flies expressing the AppA histidine acid phytase from *E. coli* had 27.82 FTU/g of phytase activity, which exceeds the 0.5-1.0 FTU/g required in animal feed. Maximum activity from AppA phytase expressed by *D. melanogaster* was observed at pH 5 and 55 °C, however, more than 50% of phytase activity was present at 25 °C and pH 2.

Here we demonstrate that insects may be suitable hosts for the heterologous expression of a microbial phytase enzyme with applications for improving animal feed nutrition and organic waste valorisation.

## Background

As global demands for meat production rise due to population growth and increased income in developing countries (Whitton et al., 2021), insects are increasingly recognised as sustainable sources of lipid and protein rich animal feed ingredients. Their ability to be reared of a range of feedstocks contributes to addressing agricultural waste management challenges and supports circular economy principles.

Easing of regulations prohibiting insect feed ingredients is also driving the insect feed industry’s growth (de Souza-Vilela et al., 2019). Insect based animal feed is approved for poultry farms and aquaculture in Australia, provided the insects are not reared on meat, manure, or postconsumer waste such as municipal biosolids (Lähteenmäki-Uutela et al., 2021). In the European Union, the use of insects for animal feed is permissible, provided they are reared exclusively on feed grade materials (Derrien, 2017). Similarly, the Association of American Feed Control Officials (AAFCO) permits the use of whole black soldier fly (*Hermetia illucens*) larvae or black soldier fly meal for use in aquaculture feed for salmonids (Lähteenmäki-Uutela et al., 2021). In South Africa, the production of insects as farm feed or pet food must undergo an application process, in which a sample of insect must be sent to a laboratory for testing (Lähteenmäki-Uutela et al., 2021). If approved, product registration is valid for 3 years, after which renewal is required (Lähteenmäki-Uutela et al., 2021). In Asia, the legislation in not well established. For example, Singapore and Thailand dominate the Asian market for insect consumables, however, the specific insect species authorised for feedstocks are yet to be defined (Deguerry et al., 2023). Similarly, South America has no official standard regarding the use of insects in animal feed (Lähteenmäki-Uutela et al., 2021).

Over 1,900 species of insects are edible. Those most frequently used commercially include yellow mealworms (*Tenebrio molitor)*, common houseflies (*Musca domestica)*, and black soldier flies (*Hermetia illucens)* (DiGiacomo and Leury, 2019). Black soldier fly (BSF) larvae are proving to be a leading choice, largely due to their ability to be reared on a broad variety of organic waste. BSF also have a favourable nutrient profile, pathogen tolerance, and are being used in an increasingly large range of animal foods, including premium cat and dog foods, aquaculture feed, small pet feed (turtle, lizard, other reptiles), poultry feed and pig feed. Genetic engineering of these edible insect species has been demonstrated (Gunther et al., 2024; Kou et al., 2023; Oppert et al., 2023; Pfitzner et al., 2024), which could increase their value as feed ingredients (Tepper et al., 2024) and accelerate the adoption of insect farming for agricultural feed ingredients globally.

Phytic acid is a major source of phosphorous that is poorly bioavailable to monogastric animals (e.g. swine, fish, and poultry), since they lack phytic acid degrading enzymes (Pramitha et al., 2021). Structurally, phytic acid is comprised of a sugar alcohol body (myo-inositol) with six phosphate groups. Phytic acid contains 12 acidic protons, which can be divided into three categories: six are strongly acidic (pK_a_ 1.5), three are weakly acidic (pK_a_ 5.7-7.6) and three are very weakly acidic (pK_a_ above 10) (Humer et al., 2015). At physiological pH (7.4) and below, phytic acid loses its protons, resulting in the negatively charged phytate anion (Costello et al., 1976).

Grains such as oats and corn, legumes, and tubers are major sources of phytic acid (Graf, 1983), with approximately 65-80% of total seed phosphorous contained in phytic acid (Gupta et al., 2015). This accounts for roughly 1-5% of total plant seed weight (Gupta et al., 2015). A diet rich in phytic acid is reflected in monogastric animal waste, with the phytate anion comprising 70% of all phosphorous excreted (Gupta et al., 2015; Lim et al., 2007). Additionally, phytate acts as an anti-nutrient. The negatively charged phosphate groups (IP_6-3_) have chelating effects on essential cationic micronutrients such as magnesium, calcium, zinc, and iron (Graf, 1983). Once bound to the phytate anion, these micronutrients are not absorbed by the digestive system, and are excreted. Moreover, diets high in phytic acid cause significant amino acid loss in livestock animals (Cowieson and Ravindran, 2007; Onyango et al., 2008). The phytate excreted in manure frequently contaminates aquatic environments by stimulating the growth of algae and cyanobacteria that can use the excess phosphorous (Correll, 1998). The resulting algae bloom formations are a major contributing factor to eutrophication (Han et al., 2012) and have severe consequences on aquatic ecosystems.

To mitigate this, phytases are frequently added to monogastric animal feed in order to increase bioavailable phosphorous in plant based feed ingredients. Phytases are enzymes that catalyse the stepwise removal of inorganic phosphate groups from the sugar alcohol body of myo-inositol (Navone et al., 2021) in phytic acid. The reaction mechanism of phytase on phytic acid can be seen in figure 1.There are a number of nutritional benefits observed when phytases are incorporated into animal feed, such as increased body weight (Wu et al., 2015), increased bone ash weight (Jones et al., 2010) and increased feed intake (Wu et al., 2015). These improved health outcomes are highly beneficial to farmers in terms of animal welfare and profitability, creating a significant market for phytase animal feed supplementation.

**Fig. 1.**
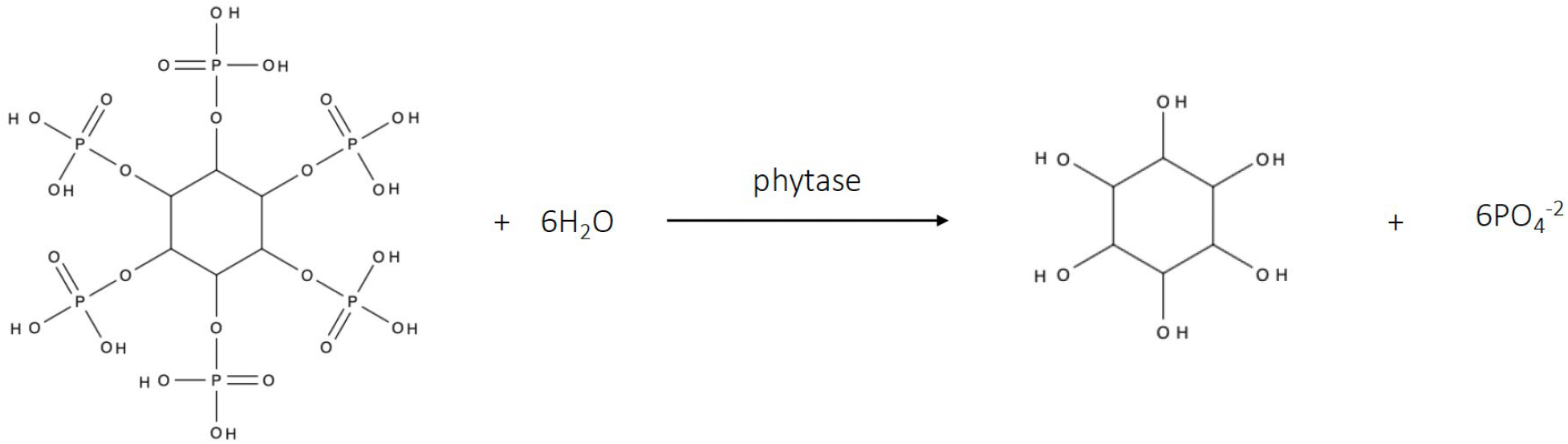
Catalytic reaction between phytic acid and phytase.

In 2023, phytase enzyme supplements had a market value of US$580 million and is expected to grow 6.6% annually (Global Market Insight, 2024). Phytases comprise over 40% of the total animal feed enzyme supplements sold – more than any other enzyme supplement (Guerrand, 2018). Phytases typically cost $0.60-$2.80 per metric tonne of feed, and this range is dependent on dose rate. Commercial phytases can be produced via microbial fermentation from bacteria or fungi, namely *Escherichia coli, Peniophora lycii, Aspergillus niger, Pichia pastoris*, and *Citrobacter brakii*.

There are four distinct classes of phytases: histidine acid phytases, β-propellor phytases, purple acid phosphatases and PTP-like phytases. Histidine acid phytases, the most prevalent phytase found in terrestrial plants, fungi, and bacterium, exhibit a notable tolerance for acidic environments (Kumar et al., 2012). β-propeller phytases are commonly found in *Bacillus* spp., and are one of the few classes of phytase that are functional in neutral to slightly alkaline environments (Greiner et al., 2007), and play a pivotal role in the phytate-phosphorus cycle in soil and water (Lim et al., 2007). Purple acid phosphatases (PAPs) are exclusive to plants, and harvest the phosphorous from the rhizosphere (Kong et al., 2018). Lastly, protein tyrosine phosphatase (PTP)-like phytases are named due to their PTP signature sequence at the active site (Gruninger et al., 2009). PTP-like enzymes are primarily found in the gut of ruminant animals and in bacteria.

Here, we engineered *Drosophila melanogaster* to express microbial, fungal and plant phytases as to determine if insects would be suitable heterologous hosts for phytase biomanufacturing. The AppA phytase expressed in transgenic *D. melanogaster* showed significant phytase activity. Our results demonstrate the first transgenic insect with functional heterologous phytase activity.

## Results

### *Drosophila melanogaster* expression plasmid design

We screened 8 different phytases; three β-propellor phytase, three purple acid phosphatase, one histidine acid phytase and one PTP like phytase (Table 1). The native signal peptide was removed and replaced with the *D. melanogaster* larval cuticle protein 9 signal peptide (Charles et al., 1998) to ensure efficient secretion by *D. melanogaster*. A short alpha tubulin promoter was selected for mid-level, ubiquitous expression in *D. melanogaster*, to mitigate the potential of toxicity.

**Table 1.** List of phytases used in this study.

| NATIVE HOST | NAME | PHYTASE CLASS | UNIPROT NO. | pH RANGE | OPTIMAL TEMPERATURE |
| --- | --- | --- | --- | --- | --- |
| *Escherichia coli* (Golovan et al., 1999) | AppA | Histidine acid phytase | P07102 | 2-5.5 | 55 °C |
| *Selenomonas ruminantium* (Yanke et al., 1999) | phyA | Beta propellor phytase | Q7WUJ1 | 4-5.5 | 50-55 °C |
| *Bacillus amyloliquefaciens* (Boukhris et al., 2015) | phy | Beta propellor phytase | O66037 | 4-10 | 70 °C |
| *Bacillus subtilis* (Tye et al., 2002) | phy | Beta propellor phytase | P42094 | 5-8 | 55 °C |
| *Glycine max*<br>(Mogal, 2017) | PAP | Purple acid phosphatase | Q09131 | 3.5-5 | 45 °C |
| *Ipomoea batatas* | PAP1 | Purple acid phosphatase | Q9SE00 | 4.5 | Unknown |
| *Arabidopsis thaliana* (Kuang et al., 2009) | PAP15 | Purple acid phosphatase | Q9SFU3 | 3.5-6 | 23-45 °C |
| *Bacillus subtilis* (Kerovuo et al., 1998) | phyC | PTP-like phytase | O31097 | 6-8 | 55 °C |

### Kinetic assays for phytases heterologously expressed in *D. melanogaster*

We screened phytase activity in transgenic adult fly lysate using a spectroscopic phytase assay kit (Megazyme, Bray, Ireland). As this method measures free phosphorous, the samples were pre-treated with an ion exchange resin to remove any free phosphorus prior to the assay. We observed clear phytase activity in the *E. coli* AppA phytase strain after a six hour incubation (Fig. 2). The remaining transgenic flies, along with the wild-type, exhibited very little or no activity (Fig. 2).

**Fig. 2.**
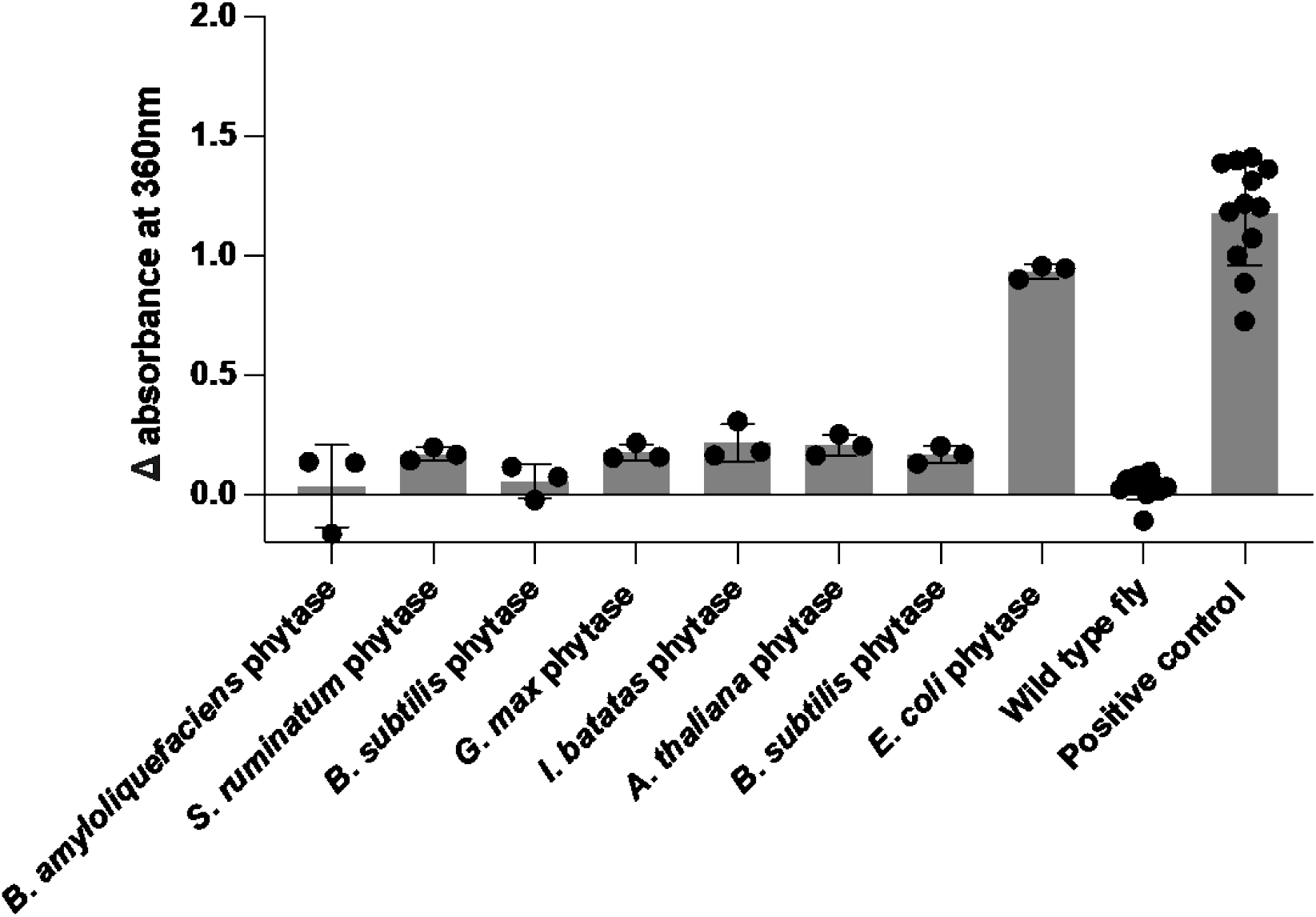
Phytase activity in transgenic flies expressing beta propellor phytase, purple acid phosphatase, PTP like phytase, and histidine acid phytase over a 6 hour incubation. n= 3-12 independent replicates with 40 adult *D. melanogaster* flies per replicate. Error bars indicate standard deviation.

### Lyophilisation assay

Only the functional *E. coli* AppA phytase was selected for further characterisation. Feed additive enzymes are often spray-dried or lyophilised and ground into a powder prior to being added to animal feed and pelleted. We tested the stability of our heterologously produced *E. coli* AppA phytase when subjected to the lyophilisation process. Phytase activity is measured in phytase units (FTU), and is defined as the amount of enzyme that liberates 1 μmol of inorganic phosphorous per minute at pH 5.5 from an excess of sodium phytate at 37 °C. Ammonium molybdate assays are the most common method to determine phytase FTU (Heinonen and Lahti, 1981; McKie and McCleary, 2016) and was used for the remaining experiments.

The average enzymatic activity of phytase expressed in lyophilised transgenic flies was determined to be 18.2 FTU/g (Fig. 3). Units were calculated using the average of three replicates at the 30 minute time point.

**Fig. 3.**
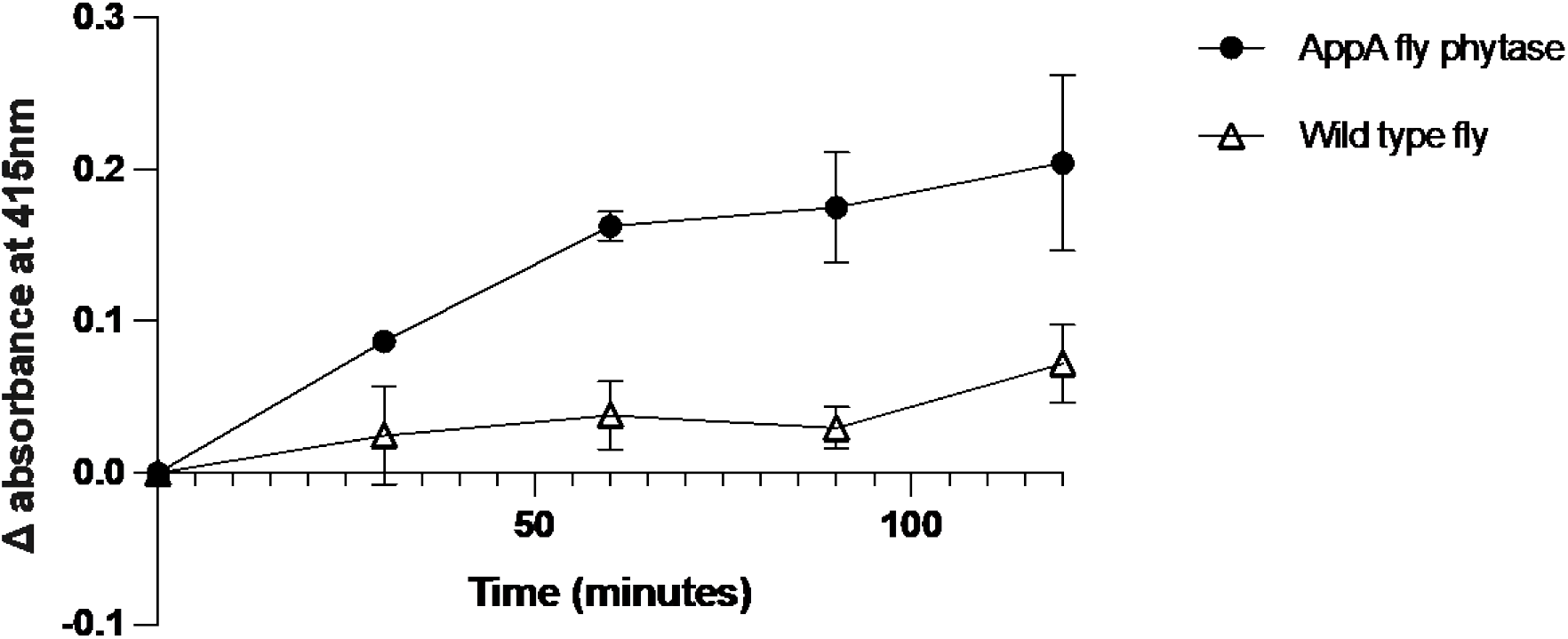
Phytase activity from lyophilised *D. melanogaster* adult flies using the ammonium molybdate method following a 120 minute incubation. n=3 biologically independent replicates. Error bars indicate standard deviation.

Figure 3 showed clear phytase activity in the transgenic *D. melanogaster* adult flies. Although minimal, the wild type flies exhibited phytase activity. The slight increase in activity observed in the wild type flies could be from endogenous phosphatases that are upregulated in the adult flies, or the expression of phosphatases from bacteria in the fly gut microbiome, both of which could hydrolyse phytic acid. According to FlyBase (FB2024_01 released February 22, 2024), there are three known gene ontologies that display phytase activity (GO:0008707, GO:0016158, and GO:0050533), and 380 phosphatase protein coding genes endogenous to adult flies. We theorised that if any of these genes were in fact hydrolysing phytic acid, they would likely be upregulated during larval and adult life stages, when dietary enzymes are required to be active for digestion, and downregulated during the pupal life stage, where digestion does not occur. To determine if this was the cause of wild type activity, the kinetic assay was repeated using *D. melanogaster* pupae rather than adults, in which dietary enzyme gene expression may be downregulated.

Figure 4 shows no phytase activity in wild type fly pupae, whilst the transgenic fly pupae retains substantial activity. The units of activity calculated for the transgenic pupae is 27.82 FTU/g .

**Fig. 4.**
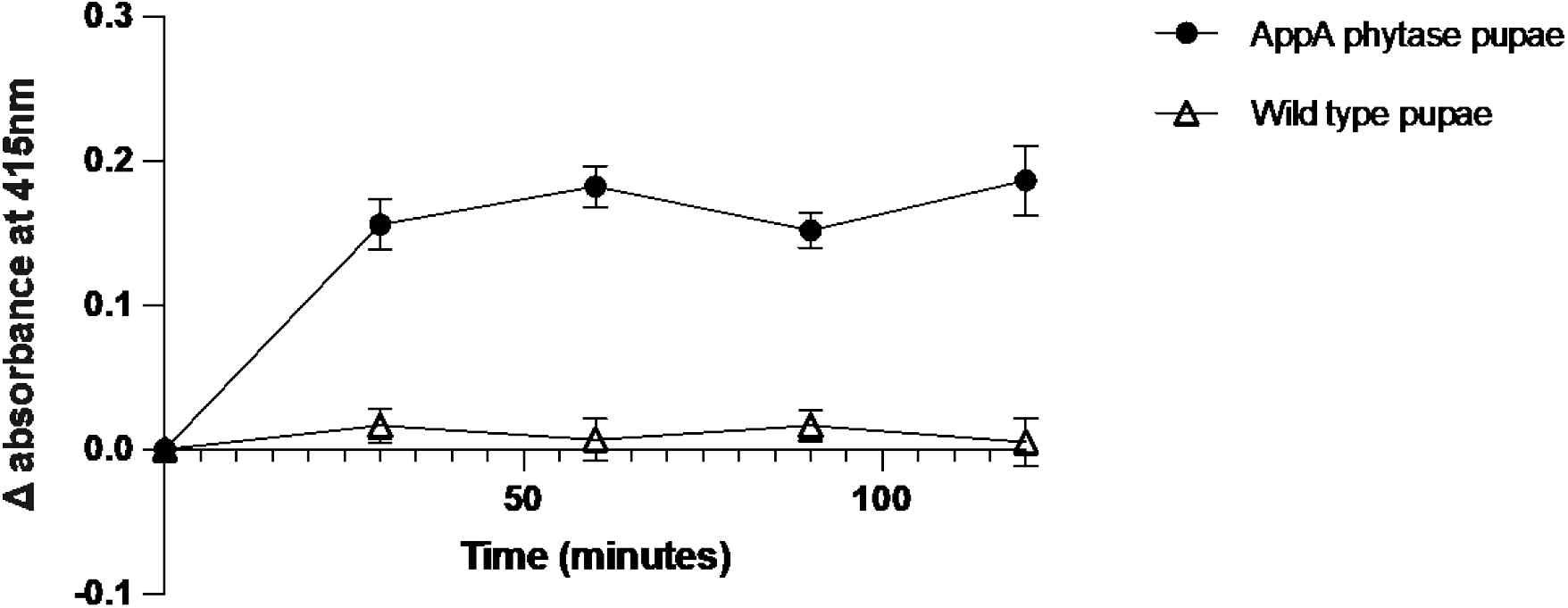
Phytase activity from lyophilised *D. melanogaster* pupae using the ammonium molybdate method following a 120 minute incubation. n=3 biologically independent replicates. Error bars indicate standard deviation.

### Phytase characterisation in *D. melanogaster*

The heterologous expression of enzymes may result in altered performance characteristics due to factors such as altered post translational modifications leading to incorrect protein folding (Shental-Bechor and Levy, 2008). To examine this, a pH and temperature curve was performed using our transgenic fly pupae.

When expressed in *D. melanogaster*, AppA phytase shows a similar pH trend compared to its endogenous expression in *E. coli*. Phytase activity is observed at pH 2-5.5, with a pH optimum of 5. At pH 2, the fly phytase retains about 60% of activity. At pH 6 and above, there is negligible phytase activity, which is consistent with the literature using bacterial and fungal hosts (Golovan et al., 1999; Navone et al., 2021).

The temperature optimum of phytase in transgenic flies is 55 °C, and a sharp decrease of activity is seen at 65 °C and above. At 25 °C, phytase activity was roughly 50% that of the optimum. Similar to the pH curve, AppA phytase activity across a range of temperatures is consistent with previous publications (Abbasi Kheirabadi et al., 2022; Wang et al., 2018).

## Discussion

In this study, we have demonstrated functional expression of *E. Coli* AppA phytase in an insect host. Moreover, the lyophilised *E. coli* AppA phytase enzyme maintained function. This is significant as many enzymes are susceptible to alpha helix decreases, beta sheet increases, and other conformation changes (Roy and Gupta, 2004) when freeze-drying, leading to decreased activity.

The commercial standard for phytases is an activity level between 500-1,000 FTU per kg of diet. The FTU determined for our transgenic flies using the ammonium molybdenum assay was 18, 210 (±1.98) FTU/kg of adult *D. melanogaster* dry weight. To produce 1kg of food supplemented with 500 FTU/kg of phytase activity, an addition of 27.5 g of our transgenic *D. melanogaster* flies would be required. Similarly, the *D. melanogaster* pupae had 27, 820 (± 13.8) FTU/kg of phytase activity, and only 17.8 g of lyophilised pupae would need to be added to 1kg of food to supply 500 FTU phytase activity. The difference in phytase activity levels reported between the adults and pupae could be due to variation in promoter activity at different developmental stages.

Several factors may account for the lack of phytase activity in the transgenic flies expressing beta propellor phytase, purple acid phosphatase, and the PTP-like phytase. For example, the glycosylation profile between bacteria, fungi, and eukaryotic systems vary dramatically (Dell et al., 2010). It is possible that the glycosylation patterns that are essential for functionality may have been compromised during the transition from the native to the insect host. Alternatively, the folding efficiency of the protein may be reduced with the loss of specific chaperone proteins from the native host. Nonetheless, the *E. coli* AppA phytase, which demonstrated functionality, will be pursued in further research using black soldier flies, including the production of thermostable phytases (Navone et al., 2021).

Here we have demonstrated that a phytase can be heterologously expressed in a transgenic insect, using the host *D. melanogaster* as proof of concept. Using insects, such as black soldier fly larvae, as livestock feed ingredients and a biomanufacturing platform will generate more value from waste streams and support more sustainable agricultural practices.

## Materials and methods

### *Drosophila melanogaster* rearing methods

Both wild type and experimental strains of *D. melanogaster* were reared in polystyrene vials (Flystuff, narrow *Drosophila* vials, #32-109RL) with a standard cornmeal diet (for 1L: 920 mL water, 67 g cornmeal, 4.6 g agar, 50 g sugar, 15.88 g yeast, 9.18 g soy flour, 60 mL of 1M propionic acid). Flies were secured in a controlled environment room at 25°C, approximately 60% humidity, and a 12 hour light/dark cycle under LED lights with a 30 minute dusk/dawn transition period.

### *Drosophila melanogaster* assembly

A *D. melanogaster* codon optimised gBlock (IDT) was synthesised consisting of the coding sequence. The plasmid consisted of a short alpha tubulin promoter, the AATCTTACAAA Kozak sequence, and SV40 terminator (Clark et al., 2022). All sequences were codon optimised for *D. melanogaster* expression using Geneious (version 2022.1.1). The gBlock (IDT) was then inserted into pMC-1-1-1 which contains a short alpha tubulin promoter and a SV40 terminator. The genes were assembled into the expression vector pMC-1-1-1 cut with NotI (Clark et al., 2022) through HiFi DNA assembly (NEBuilder HiFi DNA Assembly Master Mix NEB#E2621) into *E. coli* Stbl3 cells.

### Transformation of *Drosophila melanogaster* using microinjection

Plasmids were purified using the Monarch Plasmid MiniPrep Kit (New England Biolabs #T1010L) and sent to BestGene Inc. (Chino Hills, Ca) for embryo microinjection and subsequent PhiC31-integrase mediated transformation. Transgenic *D. melanogaster* was made via embryo microinjection and PhiC31 mediated integration into the same locus for each construct (BestGene Inc) on chromosome 2 to give the curly wing phenotype on Canton S strain (BDSC number: 64349) (Ward, 1923).

### Fly lysate preparation

To prepare the fly lysate, 40 adult flies were added for each strain in a 1.5 mL Eppendorf tube (SSIbio). All experiments with transgenic flies used strains homozygous for the phytase transgene. The Eppendorf tubes were placed in the -30°C freezer for at least 1 hour. Following the 1 hour incubation, 750 μL of 0.25M sodium acetate buffer (pH 5.5, containing 0.01% Tween 20) was added to each tube. The flies were lysed using a motorised pestle (Kimble Kontes, Pellet Pestle Motor) before incubation at 25°C for 15 minutes. The lysed flies were then centrifuged (Thermo Scientific, Heraeus Pico 17 Microcentrifuge) at 17,000 x g for 10 minutes. The supernatant was removed, taking care not to disturb the fly debris at the bottom of the tube, and placed into a fresh Eppendorf tube before centrifuging again at 17,000 x g for 10 minutes. Lastly, the supernatant was transferred to a fresh Eppendorf tube.

### UV phytase assay

The activity of the phytase enzymes in *D. melanogaster* was determined using a phytase assay kit (Megazyme; Bray, Ireland). The fly lysates for all samples were purified via ion exchange resin for the removal of free phosphate with the addition of 0.1 g of both Amberlite FPA53 OH^-^ resin (Megazyme cat. No. G-AMBOH) and Ambersep 200 H^+^ resin (Megazyme cat. No. G-AMBH) in a 1.5 mL tube. The samples were then vortexed for 30 seconds each, before being centrifuged at 3,000 x g for 10 minutes for clarification. Then, 20 μL of supernatant for each triplicate of transgenic flies, wild type flies and positive control was incubated at 40 °C for 5 minutes, and 250 μL of phytic acid was incubated simultaneously at 40 °C for 5 minutes. 20 μL of phytic acid was added to each tube containing fly lysate and vortexed thoroughly, and left at 40 °C for the appropriate time period. Following the incubation, 10 μL of stopping reagent was added to each Eppendorf tube, vortexed thoroughly, and a 10 μL aliquot of each sample was taken for the following stages.

In a UV-Star microplate (96 well, ref. no. 655801), 10 μL of sample, 60 μL of water, 20 μL of buffer, and 10 μL of MESG (2-amino-6-mercapto-7-methylpurine ribonucleoside) were added to the wells in triplicate for experimental fly strains, wild type fly strains, no fly control and a positive control (20 U/mL). Samples were mixed thoroughly by pipette, incubated for 6 hours, and the absorbance at 360nm was measured with a UV-Vis spectrophotometer (Jasco V-760 UV-Vis spectrophotometer) to obtain the background noise values (A1). Lastly, 1.5 μL of PNPase was added to each sample, and the plate was incubated at room temperature for 30 minutes before the final reading at 360nm (A2). The positive control is a purified phytase isolated from *A*.*niger*.

### Lyophilised fly preparation

Approximately 300 flies for each replicate were collected and stored in a -30 °C freezer for 7 days. The flies were then transferred to a lyophiliser (Labconco, 2.5L -50 °C series, model no.70020****) at -50 °C and 0.133mbar for 24 hours. The lyophilised flies were removed and crushed into a powder using a manual mortar and pestle. The powder was weighed and 50mg of powder was used for the experiments. The powder was hydrated using 1mL of sodium acetate buffer pH 5.5.

### Molybdenum phytase assay

A standard curve was created using 50 mM potassium phosphate stock solution and diluted to appropriate concentrations. A 5% ammonium molybdate solution was made by dissolving 2.5 g of ammonium molybdate in 50 mL water. The sulfuric acid solution was prepared by adding 7.0135 mL concentrated sulfuric acid to 12.5 mL water, and then made up to 50 mL in water. To prepare the colour reagent solution, 6.25 mL of ammonium molybdate solution and 6.25 mL sulfuric acid solution was added to 12.5 mL acetone. 20 μL of rehydrated lyophilised or lysed flies were incubated with 20 μL of 5.1 mM phytic acid sodium salt hydrate. At the appropriate time points, 20 μL of sample was thoroughly mixed with 665 μL of colour reagent solution, and absorbance was measured at 415nm. A time point 0 measurement was taken and subtracted from all following measurements to account for background absorbance.

### pH and temperature curves

For temperature curves, 50mg lyophilised flies were rehydrated with 500 μL 0.25M sodium acetate buffer pH 5.5. 20 μL of sample was mixed with 20 μL 5.1 mM phytic acid, and incubated for 90 minutes in 1.5 mL Eppendorf tubes on a heat block at varying temperatures. At the end of the incubation, 20 μL of sample was added to 665 μL of colour reagent solution, and absorbance was measured at 415nm.

For the pH curves, 50mg lyophilised flies were rehydrated in 500 μL of either 0.1M glycine-HCl buffer (pH 2-3), 0.1M acetate buffer (4-6), 0.1M Tris-HCl buffer (pH7-9) or 0.1M glycine-NaOH buffer (pH 10). Samples were incubated at 37 °C for 90 minutes, and 20 μL of sample was added to 665 μL of colour reagent solution. Absorbance was measured at 415nm.

For both temperature and pH curves, a blank was incubated alongside experimental samples consisting of 20 μL fly lysate without the phytic acid to account for any protein denaturation that may occur at extreme temperatures and pH. At the end of the incubation period, 20 μL phytic acid and 665 μL colour reagent solution was added simultaneously, and absorbance was measured at 415nm. The blank reading was subtracted from the experimental reading.

## Graph generation

Figure 1 was created using MolView 2024. Figures 2-5 was performed in GraphPad Prism version 9.4.0 (453).

**Fig. 5.**
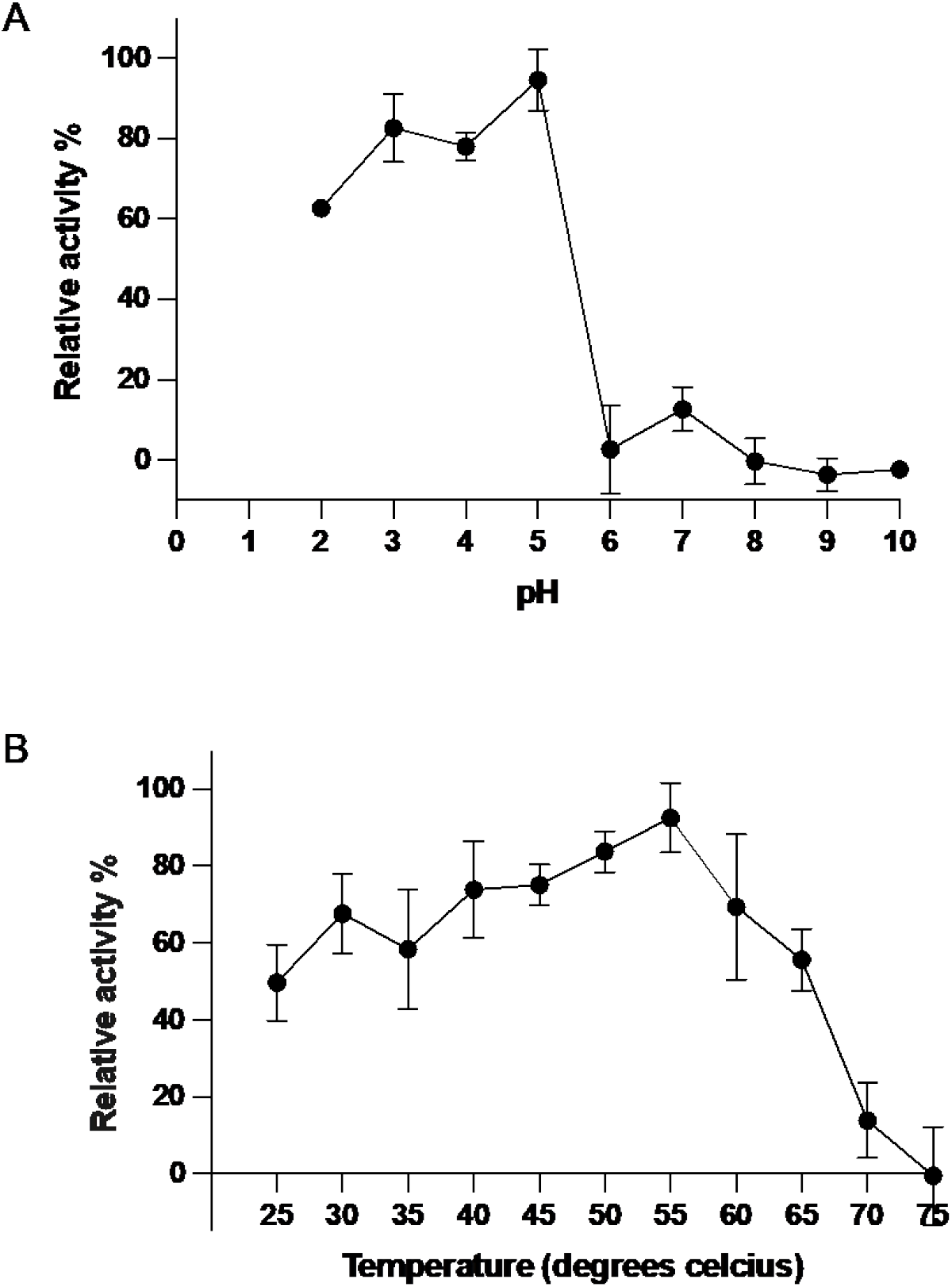
**A)** pH curve showing phytase activity of transgenic fly pupae following 90 minute incubation at 37 °C. **B)** Temperature curve showing phytase activity of transgenic fly pupae following 90 minute incubation at pH 5.5. Values reported as a percentage of the maximum phytase activity at the optimum pH/temperature observed in this assay. n=3 biologically independent replicates. Error bars indicate standard deviation.

## Data Availability

All data and calculations used in the manuscript are available in supplementary table 1.

## Acknowledgements

This research is supported by the U.S. Army International Technology Center Pacific (ITC-PAC) under Contracts FA520920P0100, FA520923C0014, and an Australian Government Research Training Program (RTP) Scholarship.

